# Temporal control of Num1 contact site formation reveals a relationship between mitochondrial division and mitochondria-plasma membrane tethering

**DOI:** 10.1101/2023.05.12.540571

**Authors:** Clare S. Harper, Jason C. Casler, Laura L. Lackner

**Affiliations:** Department of Molecular Biosciences, Northwestern University, Evanston, IL, 60208, USA

## Abstract

Mitochondrial division is critical for maintenance of mitochondrial morphology and cellular homeostasis. Previous work has suggested that the mitochondria-ER-cortex anchor (MECA), a tripartite membrane contact site between mitochondria, the ER, and the plasma membrane, is involved in mitochondrial division. However, its role is poorly understood. We developed a system to control MECA formation and depletion, which allowed us to investigate the relationship between MECA-mediated contact sites and mitochondrial division. Num1 is the protein that mediates mitochondria-ER-plasma membrane tethering at MECA sites. Using both rapamycin-inducible dimerization and auxin-inducible degradation components coupled with Num1, we developed systems to temporally control the formation and depletion of the native contact site. Additionally, we designed a regulatable Num1-independant mitochondria-PM tether. We found that mitochondria-PM tethering alone is not sufficient to rescue mitochondrial division and that a specific feature of Num1-mediated tethering is required. This study demonstrates the utility of systems that regulate contact site formation and depletion in studying the biological functions of membrane contact sites.

## INTRODUCTION

Having everything in the right place at the right time is vital for the complex coordination of cellular processes. Organelles are no exception to this, as the distribution of organelles is critical for cell health. For example, in budding yeast, several organelles need to be trafficked into the growing daughter cell and properly positioned in both the mother and daughter cell before cell division in order to ensure cell viability (Fagarasanu and Rachubinski, 2007; Eves *et al*., 2012; Knoblach and Rachubinski, 2015; Li *et al*., 2021). Organelle distribution is controlled by trafficking via motor proteins, organelle biogenesis, dynamics such as fission and fusion, and tethering to other membranes. Tethering takes place at membrane contact sites (MCSs), or areas of close apposition between membranes without fusion (Scorrano *et al*., 2019). MCSs are mediated by molecules that tether organelle membranes together through protein-protein or protein-lipid interactions. Sometimes tethering is brief and dynamic, such as with ER-Golgi contacts (David *et al*., 2021). In other cases, MCSs are stable and robust. One such example is the *S. cerevisiae* mitochondria-ER-cortex anchor (MECA), which forms a tripartite membrane tether between mitochondria, the endoplasmic reticulum (ER), and the plasma membrane (PM) that is stable throughout the cell cycle (Cerveny *et al*., 2007; Tang *et al*., 2012; Klecker *et al*., 2013; Lackner *et al*., 2013).

MECA is composed of Num1, a 313 kDa protein that interacts directly with the mitochondrial outer membrane through its N-terminal coiled-coil (CC) domain (Tang *et al*., 2012; Lackner *et al*., 2013; Ping *et al*., 2016), and the PM through its C-terminal pleckstrin homology (PH) domain, which binds with high specificity to the PM-enriched phosphoinositide phosphatidylinositol 4,5-bisphosphate [PI(4,5)P2] (Yu *et al*., 2004; Tang *et al*., 2009). The association of Num1 with the ER is thought to be mediated by an interaction between Num1 and the integral ER membrane protein Scs2 (Chao *et al*., 2014; Omer *et al*., 2018) (Figure 1A). The association between Num1 and mitochondria drives the assembly of Num1 into stable clusters that tether the three membranes together (Kraft and Lackner, 2017). When mitochondria are trafficked into the growing daughter cell, MECA-mediated mitochondrial tethering helps retain mitochondria in the mother cell (Cerveny *et al*., 2007; Klecker *et al*., 2013; Lackner *et al*., 2013). Following mitochondrial inheritance, Num1 clusters form and tether mitochondria in the bud, ensuring mitochondria are retained in the bud during cell division (Kraft and Lackner, 2017).

**Figure 1:**
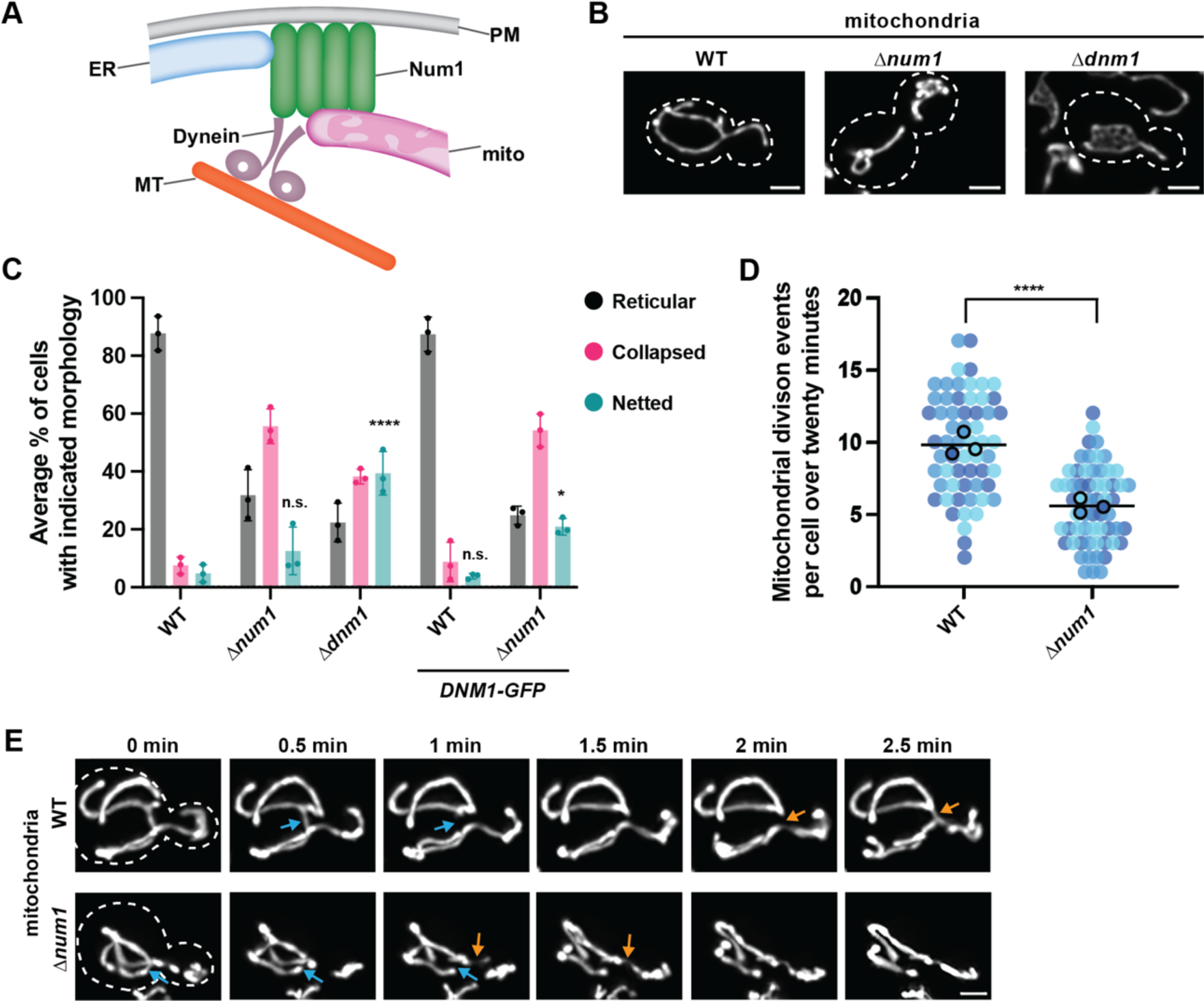
Num1 facilitates mitochondrial division. (A) Schematic of the mitochondria-ER-cortex-anchor (MECA) contact site. (B) Wild type (WT), Δ*num1*, and Δ*dnm1* cells expressing mito-Red were analyzed by fluorescence microscopy. Cells were grown in SCD at 30°C. Whole-cell maximum projections and single slice images are shown. The cell cortex is outlined with a white dashed line. Bar, 2 µm. (C) Quantification of mitochondrial morphology (either reticular, collapsed, or netted) for WT, Δ*num1*, Δ*dnm1, DNM1-GFP* and Δ*num1 DNM1-GFP* cells is shown as the mean ± SD. Each black dot represents the average for one biological replicate, with >100 cells quantified per replicate. Three biological replicates are shown. Representative images of each morphology category are shown to the right. *p* values are in comparison to wild type. * *p* < 0.05; **** *p* < 0.0001; n.s., not significant (Mann-Whitney test). (D) Quantification of mitochondrial division events per cell over a twenty-minute time-lapse movie. Each dot represents a single cell. There are 20 cells per biological replicate, and the three biological replicates are represented in different colors. The black line denotes the grand mean. *p* values are in comparison to wild type. **** p < 0.0001 (Mann-Whitney test). (E) Examples of fusion and division events for WT and Δ*num1* cells expressing mito-Red are shown. Whole cell maximum projections. Timepoints are shown in minutes. Blue arrows indicate mitochondrial division events and orange arrows indicate mitochondrial fusion events. Bar, 2 µm.

Similar to many MCS proteins, Num1 is multifunctional and has additional roles in the cell besides tethering organelles (Eisenberg-Bord *et al*., 2016; Lackner, 2019; Scorrano *et al*., 2019; Harper *et al*., 2020; Prinz *et al*., 2020). In addition to maintaining mitochondrial distribution during inheritance, Num1 is required for the function of dynein in nuclear inheritance. Dynein is trafficked to the plus ends of astral microtubules and offloaded onto Num1 clusters at the cortex. Num1-anchored dynein then captures and walks along astral microtubules, generating the force to align the mitotic spindle for nuclear inheritance (Kormanec *et al*., 1991; Eshel *et al*., 1993; Carminati and Stearns, 1997; Adames and Cooper, 2000; Heil-Chapdelaine *et al*., 2000; Farkasovsky and Küntzel, 2001; Lammers and Markus, 2015). Interestingly, Num1-mediated dynein anchoring is also important for mitochondrial function; when dynein is unable to anchor at cortical Num1 clusters, defects in mitochondrial respiration are observed (White *et al*., 2022). Thus, although MCS proteins have roles in seemingly distinct pathways, these roles may be interrelated in order to coordinate cellular processes.

In addition to its role in mitochondrial tethering and dynein anchoring, Num1 has also been implicated in mitochondrial division (Cerveny *et al*., 2007). Mitochondrial division and the opposing activity of mitochondrial fusion are critical for maintaining the shape and health of the mitochondrial network and the integrity of mitochondrial DNA (mtDNA) (Hoppins *et al*., 2007; Nunnari and Suomalainen, 2012; Osman *et al*., 2015). The mechanisms of mitochondrial division and fusion are conserved from yeast to metazoa (Hoppins *et al*., 2007). In yeast, the dynamin-related GTPase Dnm1 drives mitochondrial division. Dnm1 is recruited to the mitochondrial surface by the adaptor protein Mdv1, which binds to the outer mitochondrial membrane protein Fis1 (Mozdy *et al*., 2000; Tieu and Nunnari, 2000; Cerveny *et al*., 2001; Tieu *et al*., 2002; Cerveny and Jensen, 2003). Mdv1 subsequently helps drive the assembly of Dnm1 into a helical structure that wraps around the organelle, causing constriction and ultimately scission of the mitochondrial membranes (Naylor *et al*., 2006; Lackner *et al*., 2009; Mears *et al*., 2011). Num1 was identified in a genetic screen as a putative Dnm1-interacting protein and was suggested to be critical for mitochondrial division (Cerveny *et al*., 2007). A subsequent study indicated that Num1 is not required for mitochondrial division, but that the rates of mitochondrial division are reduced in cells lacking Num1 (Lackner *et al*., 2013). However, the mechanism by which Num1 impacts mitochondrial division is unclear. One hypothesis is that Num1-mediated tethering of mitochondria creates tension across the mitochondrial membrane as it is stretched between anchor points (Schauss and McBride, 2007; Lackner and Nunnari, 2009; Hammermeister *et al*., 2010). Several studies suggest that membrane tension is important for the ability of dynamin-related proteins to mediate membrane scission (Roux *et al*., 2006; Mahecic *et al*., 2021). Another hypothesis is that Num1 plays an active role in division, perhaps promoting or regulating Dnm1 function. Interestingly, a population of Dnm1 colocalizes with Num1 at the cell cortex (Cerveny *et al*., 2007; Lackner *et al*., 2013); however, the reason for this colocalization and the role of this population of Dnm1 remains unclear.

Here, we sought to further clarify the role of Num1 in mitochondrial division and address the relationship between Num1-mediated mitochondrial tethering and mitochondrial division. To this end, we developed a system to temporally control the formation and depletion of mitochondria-PM contact sites using regulatable versions of the native Num1 tether as well as an artificial tether. Using these tools, we found that mitochondrial division rates are reduced shortly after Num1-mediated mitochondrial tethering to the cell cortex is lost and are rescued shortly after tethering is induced. Surprisingly, using an artificial, Num1-independent inducible tether, we found that mitochondria-PM tethering alone is not sufficient to rescue mitochondrial division and that a specific feature or function of Num1-mediated tethering is required. Tools for the temporal control of MCS formation and depletion using native and artificial membrane tethers, like those presented in this study, are useful to determine the distinct features and activities of MCS proteins that contribute to the biological functions of the MCS, as well as the kinetics of the events that occur downstream of MCS formation.

## RESULTS

### Loss of Num1 results in disrupted mitochondrial division

Num1 has been suggested to be required for mitochondrial division; cells lacking Num1 have been reported to have net-like mitochondrial networks and very few to no observable mitochondrial division events (Cerveny *et al*., 2007; Hammermeister *et al*., 2010). In contrast, our previous work suggests that Δ*dnm1* and Δ*num1* cells have distinct defects in mitochondrial morphology and, while mitochondrial division may be attenuated in the absence of Num1, Num1 does not play a direct role in mitochondrial division (Lackner *et al*., 2013). To clarify the role of Num1 in mitochondrial division, we re-examined the morphology of mitochondria in wild type, Δ*dnm1*, and Δ*num1* cells expressing mitochondrial matrix-targeted dsRED (mito-Red) using high-resolution microscopy (Figure 1B). Mitochondrial morphology was analyzed on a per cell basis and the networks were classified as follows: reticular (mitochondrial tubules interlaced yet spread apart, with few junctions), collapsed (mitochondrial tubules hard to distinguish from each other because they are entangled or clumped together), or netted (several closely apposed junctions in the mitochondrial network causing fenestrations) (Figure 1C and S1). Cells lacking Num1 predominantly displayed a collapsed mitochondrial network, clearly distinct from the hyperfused, netted mitochondrial networks observed in the majority of Δ*dnm1* cells (Figure 1B and C). Through rigorous quantification of high-resolution time-lapse images, we observed that the number of division events over twenty minutes is reduced in Δ*num1* cells compared to wild type cells (Figure 1D and E). These results are consistent with our previous findings that mitochondria in Δ*num1* cells and Δ*dnm1* cells have distinct morphological defects and that mitochondrial division occurs at a reduced frequency in cells lacking Num1 (Lackner *et al*., 2013).

We next sought to examine Dnm1-marked sites of mitochondrial division in Δ*num1* cells. We expressed Dnm1 as a yEGFP (GFP) fusion from its endogenous locus in Δ*num1* cells expressing mito-Red. Surprisingly, in comparison to Δ*num1* cells expressing untagged Dnm1, we observed an increase in the percentage of Δ*num1 DNM1-GFP* cells that display a hyperfused, netted mitochondrial network (12% and 21% of cells, respectively; Figure 1C). The netted phenotype was not as severe as a complete disruption of division, where 40% of Δ*dnm1* cells have a netted mitochondrial network (Figure 1C). In our previous work, we did not observe netted mitochondrial networks in Δ*num1* cells expressing GFP-tagged Dnm1 (Lackner *et al*., 2013). However, in that study, we expressed GFP-Dnm1 from a plasmid in addition to untagged Dnm1 from its endogenous locus. We replicated the results of our previous study, finding that a very small percentage of Δ*num1* cells (1.6% of cells) exhibited netted mitochondrial networks when GFP-Dnm1 was expressed from a plasmid (Figure S2C-D). These results suggest that the increase in cells with netted mitochondrial networks is due to the expression of Dnm1-GFP from the endogenous locus. We hypothesize that the C-terminal GFP tag on Dnm1 partially compromises its function, creating a hypomorphic allele. In a wild type background, the hypomorphic phenotype of Dnm1-GFP is not evident (Figure 1C). The phenotype only becomes apparent in the absence of Num1, when mitochondrial division is already compromised. The hypomorphic Dnm1 allele fortuitously provides us with a way to further enhance the mitochondrial division defect in Δ*num1* cells and more clearly assess the role of Num1 in mitochondrial division.

### Characterizing a system to control the formation and depletion of mitochondria-ER-PM contact sites

To further assess the role of Num1 in mitochondrial division, we sought to develop a system in which we could control the formation and depletion of Num1-mediated contact sites. Such a system would allow us to ask if and how quickly mitochondrial division could be restored following the induction of Num1-mediated mitochondrial tethering. We engineered an inducible Num1-mediated contact site using the rapamycin-inducible dimerization system. Since rapamycin inhibits the TOR pathway in yeast, we used a rapamycin resistant strain background, which harbors the following mutations: Δ*fpr1 tor1-1* (Gruber *et al*., 2006; Haruki *et al*., 2008). We first confirmed that deletion of *NUM1* in the rapamycin resistant background led to mitochondrial morphology and division defects similar to those observed in Δ*num1* cells (Figure S2A-B and Figure 1B-D, respectively). Next, we appended the rapamycin-inducible dimerization components FRB-GFP and FKBP12 to Num1ΔPH (amino acids 1-2562) and the Num1 PH domain (Num1PH, amino acids 2563-2748), respectively (Figure 2A). Num1ΔPH-FRB-GFP was expressed from the endogenous *NUM1* locus and *FKBP12-NUM1PH* was integrated into the genome at the *URA* locus and expressed from the TEF promoter. We refer to this strain as RID-Num1 for rapamycin-inducible dimerization of Num1. In the absence of rapamycin (uninduced RID-Num1 or uRID-Num1), Num1ΔPH-FRP-GFP was diffusely distributed throughout the cytosol (Figure 2B). Following the addition of rapamycin (induced RID-Num1 or iRID-Num1), cortical, mitochondria-associated clusters of Num1ΔPH-FRB-GFP were observed (Figure 2B, Figure S3A, Video 1). We refer to these clusters as RID-Num1 clusters. We found that expression of FKBP12-Num1PH from the mid-strength TEF promoter best recapitulated wild type Num1 cluster formation, mitochondrial tethering, and mitochondrial function when compared to expression from the comparatively weaker and stronger ADH and GPD promoters, respectively (Figure S3B-C). Formation of RID-Num1 clusters occurred rapidly following rapamycin addition; within 5 minutes the number of RID-Num1 clusters formed was comparable to the number of wild type Num1 clusters (Figure 2B and C). The RID-Num1 clusters robustly tethered mitochondria to the PM (Figure 2D, Figure S3A, Video 1). Thus, the RID-Num1 system provides a way to rapidly induce Num1-mediated mitochondria-PM tethering.

**Figure 2:**
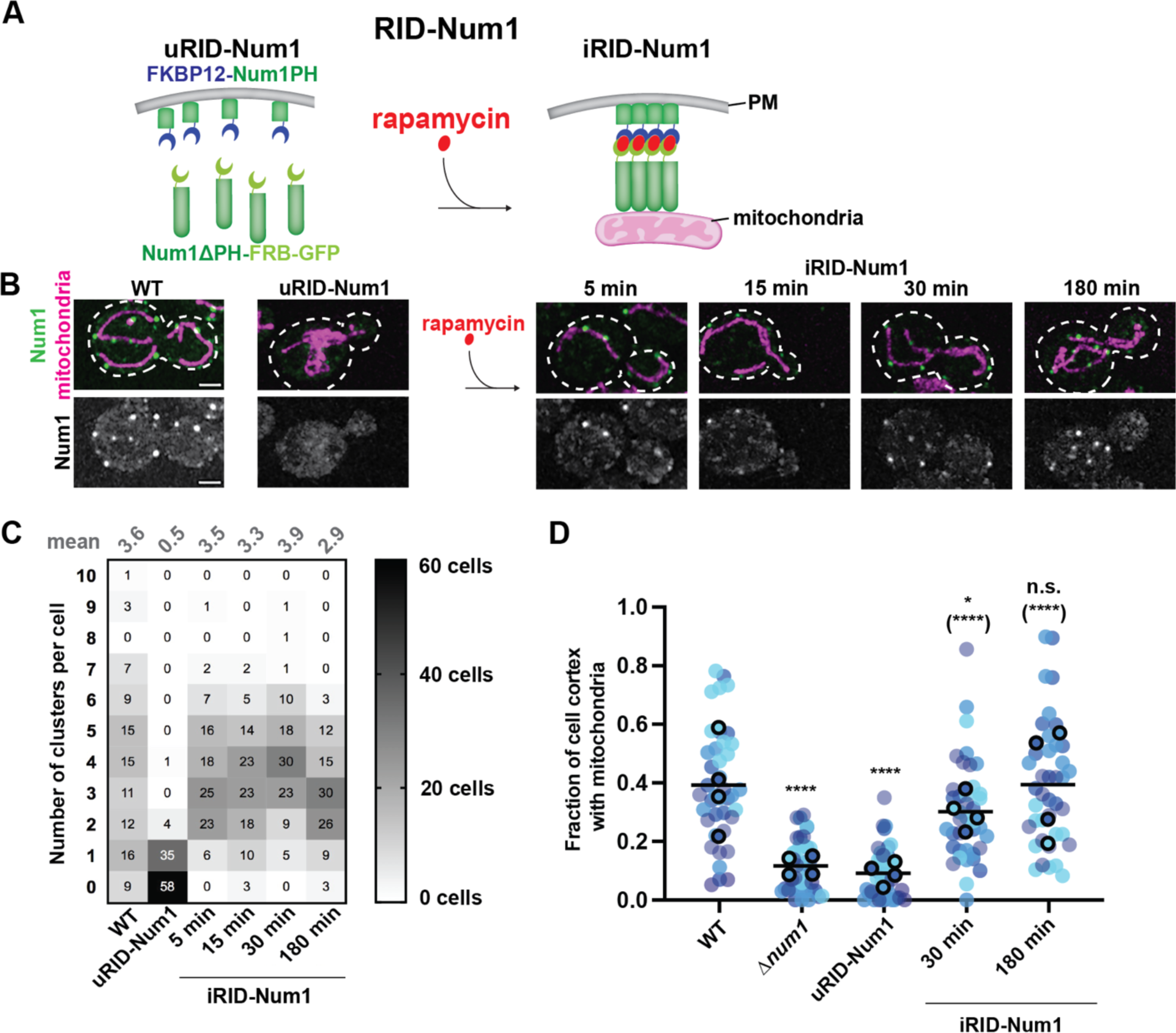
A regulatable system to control the formation of Num1 clusters. (A) Schematic depicting the rapamycin-induced dimerization (RID-Num1) system to inducibly form Num1-mediated contact sites. The left schematic represents the components of the uninduced RID-Num1 system before rapamycin addition, and the right schematic represents the RID-Num1 Num1 contact sites that form after dimerization is induced by rapamycin addition. uRID-Num1, uninduced RID-Num1; iRID-Num1, induced RID-Num1; PM, plasma membrane. (B) *NUM1-GFP* and RID-Num1 cells expressing mito-Red were analyzed by fluorescence microscopy. Whole-cell maximum projections are shown. The far left is a cell expressing *NUM1-GFP* and mito-Red with no rapamycin added. RID-Num1 cells are shown without (uRID-Num1) and with (iRID-Num1) rapamycin. A different cell is shown for each timepoint. The top panel shows a merge of the GFP and mito-Red signals and the bottom panel shows the GFP signal in greyscale. Bar, 2 µm. (C) Quantification of the number of RID-Num1 clusters per cell for WT cells expressing *NUM1-GFP* and RID-Num1 cells. RID-Num1 cells were analyzed at the indicated times after rapamycin addition. n = 100 cells over three image replicates. (D) Quantification of the fraction of the cell cortex at mid-cell occupied by mitochondria for WT, Δ*num1*, uRID-Num1, and iRID-Num1 cells. Each dot represents a single cell and each biological replicate is represented by a different color. n = 30 cells. *p* values are in comparison to WT, *p* values in parenthesis are in comparison to uRID-Num1. * *p* < 0.05; **** *p* < 0.0001; n.s. not significant (Mann-Whitney test).

Next, we built upon the RID-Num1 system to make it reversible. We added an auxin-inducible degron (AID) tag to FKBP12-Num1PH, constructing FKBP12-AID-Num1PH, which we expressed in the rapamycin resistant strain along with Num1ΔPH-FRB-GFP and Tir1. Tir1 is a plant specific F-box protein that binds the yeast SCF (Skp1, Cullen, F-box) complex. In the presence of auxin, Tir1 recruits the SCF complex to AID tagged proteins, which are subsequently ubiquitinated and targeted for proteasomal degradation (Nishimura *et al*., 2009; Morawska and Ulrich, 2013). We refer to this modified RID-Num1 system as RID^AID^-Num1 (Figure 3A). Similar to the RID-Num1 system, we observed RID^AID^-Num1 clusters form at the PM following rapamycin addition, and these clusters tethered mitochondria (Figure 3B). The subsequent addition of auxin to induced RID^AID^-Num1 (iRID^AID^-Num1) cells resulted in the depletion of RID^AID^-Num1 clusters (dRID^AID^-Num1), with the maximal depletion occurring between 2 and 2.5 hours after auxin treatment (Figure 3B and C). Though this is longer than most AID-tagged protein depletions, it was not unexpected given that Num1 clusters are very stable and display limited exchange with the non-clustered pool of Num1 (Kraft and Lackner, 2017). Immunoblotting revealed that not only was FKBP12-AID-Num1PH degraded after auxin addition, but Num1ΔPH-FRB-GFP was also degraded, emphasizing the stability of Num1 clusters and strength of the rapamycin-induced dimerization interaction (Figure 3E, full blots in Figure S4A-B). As expected, in uninduced RID^AID^-Num1 cells (uRID^AID^-Num1), which were never exposed to rapamycin, the addition of auxin resulted in the degradation of FKBP12-AID-Num1PH, but not Num1ΔPH-FRB-GFP (Figure S4C). Consistent with Num1 cluster depletion in dRID^AID^-Num1 cells, mitochondria-PM tethering was also reduced in these cells, resulting in collapsed mitochondrial networks similar to those observed in Δ*num1* cells (Figure 3B and D). Thus, by adding the AID tag to the RID-Num1 system, we have engineered a strain that allows us to induce and subsequently deplete Num1-mediated mitochondria-PM tethering and examine the downstream consequences.

**Figure 3:**
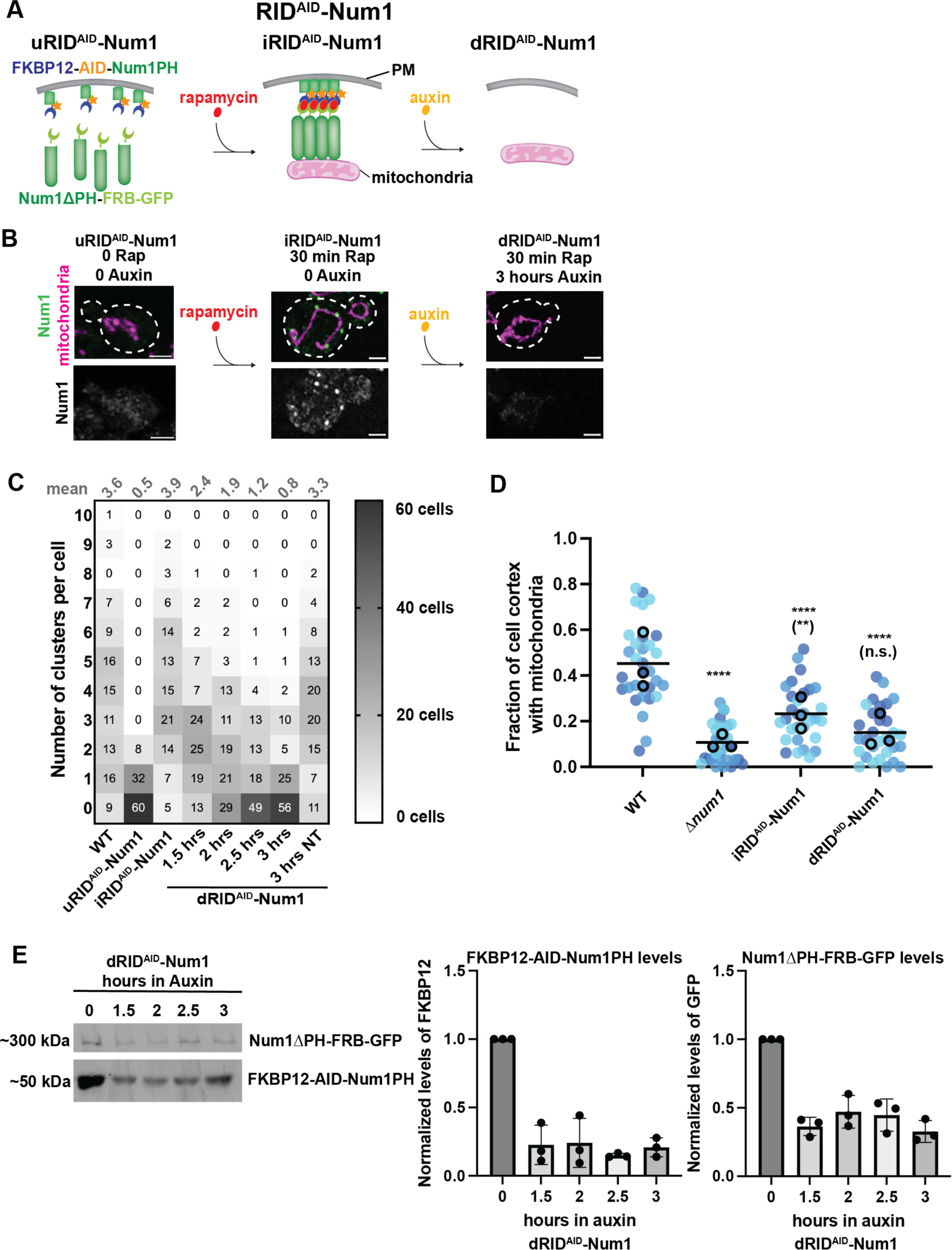
A regulatable system to deplete Num1 clusters. (A) Schematic depicting the rapamycin-induced dimerization auxin-inducible degradation (RID^AID^-Num1) system to inducibly form and deplete Num1-mediated contact sites. The left schematic represents the components of RID^AID^-Num1 before rapamycin addition, the middle schematic represents the Num1 contact sites that form after dimerization is induced by rapamycin addition, and the right schematic represents the system after addition of auxin and degradation of Num1 components. (B) RID^AID^-Num1 cells expressing mito-Red were analyzed by fluorescence microscopy. Whole-cell maximum projections are shown. RID^AID^-Num1 cells without any treatment (uRID^AID^-Num1), with rapamycin added (iRID^AID^-Num1), and with rapamycin and auxin added (dRID^AID^-Num1) are shown. A different cell is shown for each condition. The top panel shows a merge of the GFP and mito-Red signals and the bottom panel shows the GFP signal in greyscale. Bar, 2 µm. (C) Quantification of the number of Num1 RID^AID^-Num1 clusters per cell for WT cells expressing *NUM1-GFP* and RID^AID^-Num1 cells. iRID^AID^-Num1 cells were incubated with rapamycin for 30 minutes. dRID^AID^-Num1 cells were incubated with rapamycin for 30 minutes, washed with media, and then incubated with auxin for the indicated times, in hours. The “3 hrs NT” sample was treated with rapamycin for 30 minutes, washed with media, and then incubated with no treatment for three hours. n = 100 cells over three image replicates. (D) Quantification of the fraction of the cell cortex at mid-cell occupied by mitochondria for iRID^AID^-Num1 and dRID^AID^-Num1 cells. Data for WT and Δ*num1* are duplicated from Figure 2D. Each dot represents a single cell and each biological replicate is represented by a different color. n = 30 cells. *p* values are in comparison to WT, *p* values in parenthesis are in comparison to Δ*num1*. ** *p* < 0.01; **** *p* < 0.0001; n.s. not significant (Mann-Whitney test). (E) Western blot of RID^AID^-Num1 components (detecting Num1ΔPH-FRB-GFP with an α-GFP antibody and FKBP12-Num1PH with an α-FKBP12 antibody). RID^AID^-Num1 cells were incubated with rapamycin for 30 minutes, auxin was then added, and the cells were analyzed at the indicated times after auxin addition. Quantification of normalized protein levels is shown as the mean ± SD, n=3 independent experiments. Total protein stain was used as a loading control (see Figure S4B) as described in the Materials and Methods.

### Synthetic, regulatable Num1 contact sites behave like native Num1 contact sites

To fully validate the contact sites formed in the RID-Num1 and RID^AID^-Num1 systems, it was important to establish that they function similarly to native Num1 contact sites, which have roles beyond mitochondria-PM tethering. Native Num1 clusters associate with the ER, in addition to mitochondria and the PM. We verified that RID-Num1 and RID^AID^-Num1 clusters associate with the ER by live cell microscopy. We examined the localization of RID-Num1 and RID^AID^-Num1 clusters relative to the ER, and we found that 96.8% RID-Num1 clusters and 95% RID^AID^-Num1 clusters (n>175 clusters) were colocalized with the ER (Figure 4A). Because the cortical ER covers a significant portion of the cell cortex, we also examined the localization of RID-Num1 and RID^AID^-Num1 clusters relative to the ER in Δ*lnp1* cells, in which the distribution of the cortical ER is disrupted, resulting in large ER-free regions at the cell cortex (Chen *et al*., 2012). In the Δ*lnp1* background, the colocalization of RID-Num1 and RID^AID^-Num1 clusters with ER was still apparent (96.6% and 95.7%, respectively; n>175 clusters) (Figure 4A).

**Figure 4:**
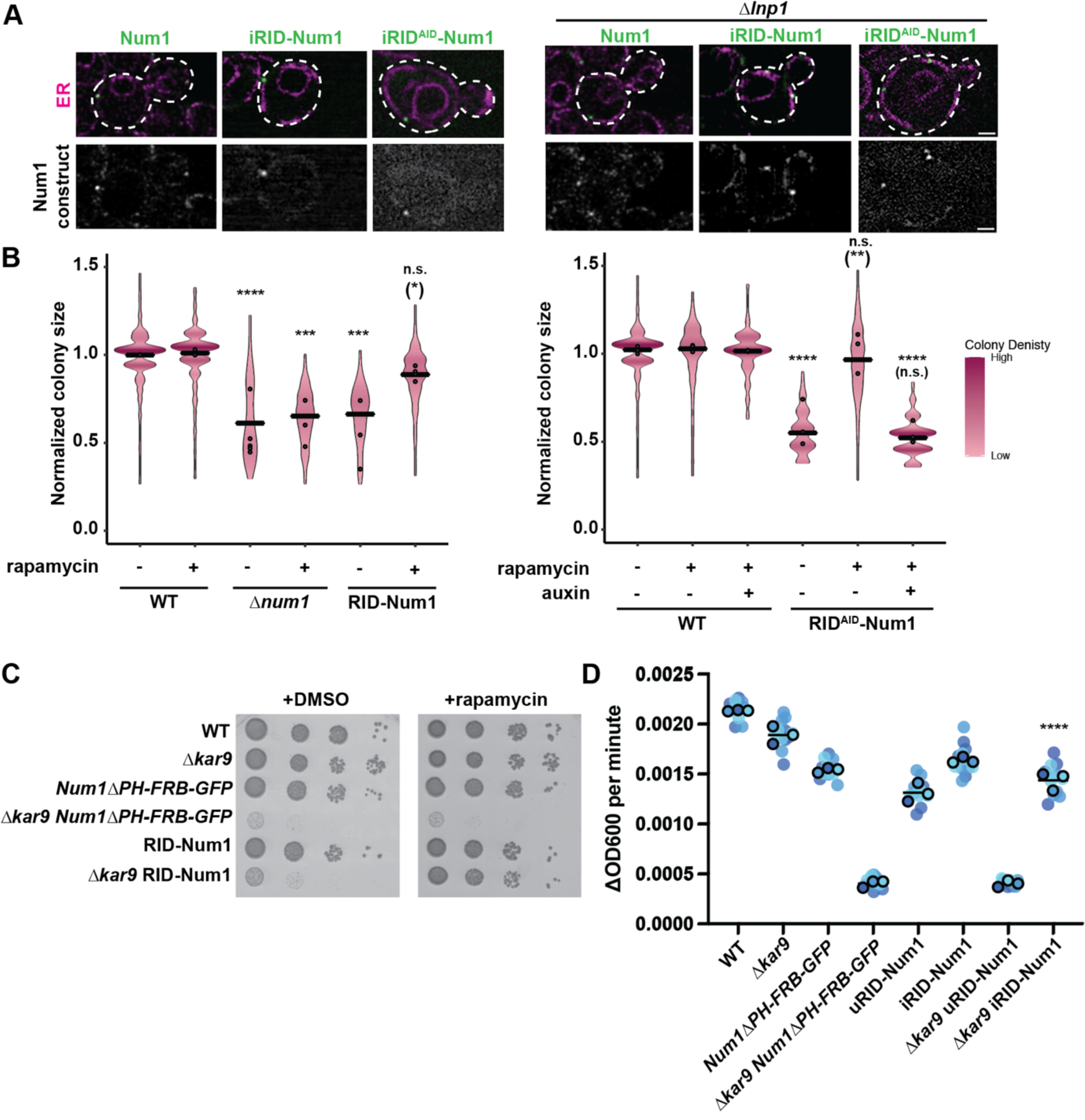
RID-Num1 and RID^AID^-Num1 systems behave like wild type Num1-mediated contact sites. (A) *NUM1-GFP*, RID-Num1, and RID^AID^-Num1 cells expressing ER-Red in both wild type and Δ*lnp1* backgrounds were analyzed by fluorescence microscopy. Single focal planes are shown. RID-Num1 and RID^AID^-Num1 cells were imaged 30 minutes after rapamycin addition. Bar, 2 µm. (B) Quantification of WT, Δ*num1*, RID-Num1, and RID^AID^-Num1 colony size, with cells grown at 35°C in respiratory growth conditions (YPEG solid media) for 5 days. DMSO (1 µM), rapamycin (1 µM), or auxin (1 mM) was added to plates, as indicated. The graph is a violin plot of the radius (in pixels) of colonies normalized to the mean radius of WT colonies for the respective experiment. The black line denotes the grand mean of at least three independent experiments and dots depict the mean of each experiment. n>300 colonies. *p* values are in comparison to WT, *p* values in parenthesis are in comparison to RID-Num1 without rapamycin and RID^AID^-Num1 without rapamycin or auxin, respectively. ** *p* < 0.01; *** *p* < 0.001; **** *p* < 0.0001; n.s. not significant (Mann-Whitney test). (C) Spot test growth assay of WT, *Δkar9, NUM1ΔPH-FRB-GFP, Δkar9 NUM1ΔPH-FRB-GFP,* RID-Num1, and *Δkar9* RID-Num1 strains in fermentative growth conditions (YPD solid media) at 30°C for 1 day. DMSO or 1 µM rapamycin were added to plates, as indicated. (D) The change in OD600 per minute for the linear portion of the growth curves obtained using a microplate reader for WT, *Δkar9, NUM1ΔPH-FRB-GFP, Δkar9 NUM1ΔPH-FRB-GFP,* RID-Num1, and *Δkar9* RID-Num1 strains in fermentative growth conditions (YPD media) at 30°C for 1 day. Dots represent biological replicates, each composed of three to four technical replicates; mean ± SD is shown. *p* values are in comparison to Δ*kar9* uRID-Num1. **** *p* < 0.0001 (Mann-Whitney test).

Additionally, we tested the ability of RID-Num1 and RID^AID^-Num1 clusters to support mitochondrial and dynein function, which are both negatively affected by disruption of Num1 contact sites. In the absence of rapamycin, both uRID-Num1 and uRID^AID^-Num1 cells exhibited an impairment in respiratory growth similar to that of Δ*num1* cells, as assessed quantitatively by colony size, indicative of a defect in mitochondrial function. In the presence of rapamycin, which induced RID-Num1 and RID^AID^-Num1 cluster formation and restored mitochondrial tethering (Figure 2A-D and 3A-D), respiratory growth was rescued (Figure 4B). The subsequent addition of auxin to iRID^AID^-Num1 cells resulted in a reduction in respiratory growth to a level similar to that observed for Δ*num1* cells, consistent with depletion of the RID^AID^-Num1 clusters.

Our previous work found that Num1-mediated cortical dynein anchoring is important for normal growth under respiratory conditions (White *et al*., 2022). The increased respiratory growth observed for iRID-Num1 cells in comparison to the uninduced state suggests that RID-Num1 clusters are able to anchor dynein. To further assess the ability of RID-Num1 clusters to anchor dynein as well as support dynein function in nuclear migration, we examined the growth of RID-Num1 cells in the absence of Kar9. Kar9 is required for a partially redundant nuclear inheritance pathway. Loss of both the dynein and Kar9 pathways for nuclear inheritance results in a severe growth defect (Miller and Rose, 1998). In a Δ*kar9* background, uRID-Num1 cells exhibited a severe growth defect, similar to Δ*kar9 NUM1ΔPH-FRB-GFP* cells (Figure 4C-D). Following the addition of rapamycin, growth of the Δ*kar9* iRID-Num1 cells was significantly increased to near wild type levels (Figure 4C-D). These results indicate that the RID-Num1 clusters support dynein function in nuclear inheritance. Finally, we found that there is a stable cortical population of Dnm1 in iRID-Num1 cells, similar to wild type (Figure S5, Video 2). Thus, the inducible and reversible synthetic Num1 tethers recapitulate the known activities of native Num1.

### Tethering mitochondria to the plasma membrane with regulatable Num1 contact sites restores mitochondrial division

With the fully validated regulatable Num1 contact site systems in hand, we investigated the relationship between Num1 contact site formation/depletion and mitochondrial division. We first sought to determine if we could restore mitochondrial division rates following the formation of Num1 contact sites. To this end, we quantified the number of mitochondrial division events in uRID-Num1 and iRID-Num1 over twenty-minute time-lapse movies. The uRID-Num1 strain had decreased mitochondrial division rates, comparable to Δ*num1* cells (Figure 5A). This indicates that mitochondrial division is impaired in the absence of mitochondria-PM tethering, even when all domains of Num1 are present. Thirty minutes after rapamycin addition, mitochondrial division rates in iRID-Num1 cells approached wild type levels (Figure 5A). At longer time points post-rapamycin addition, division rates were consistently increased over uRID-Num1. These results indicate that mitochondrial division rates are restored following the formation of Num1-mediated mitochondrial tethering sites.

**Figure 5:**
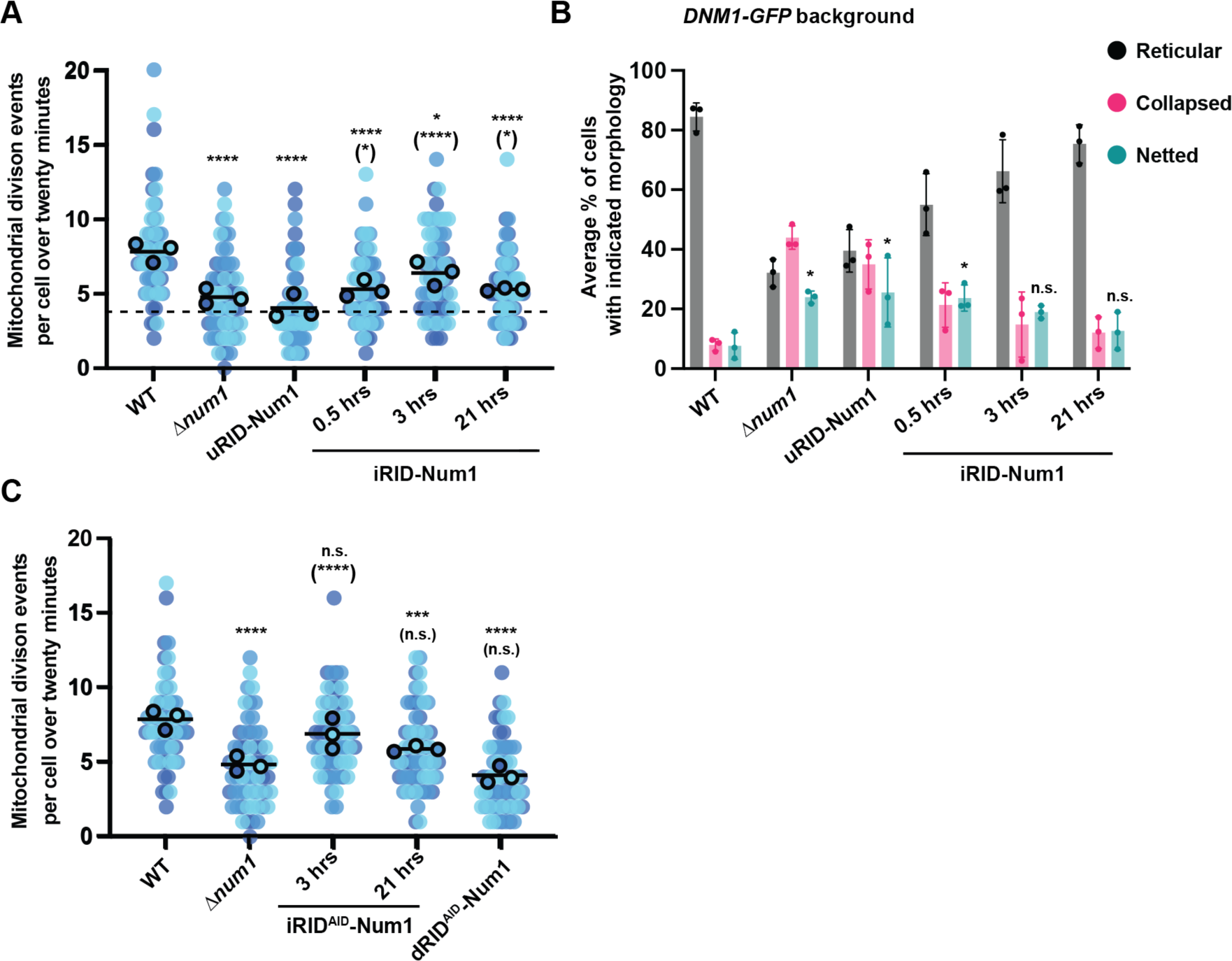
The RID-Num1 and RID^AID^-Num1 systems provide insight into the connection between Num1-mediated contact sites and mitochondrial division. (A) Quantification of mitochondrial division events per cell over a twenty-minute time-lapse movie for RID-Num1 cells treated with rapamycin for the indicated times. Each dot represents a single cell. There are 20 cells per biological replicate, and the three biological replicates are represented in different colors. The black line denotes the grand mean. *p* values are in comparison to WT, *p* values in parenthesis are in comparison to uRID-Num1. * *p* < 0.05; **** *p* < 0.0001 (Mann-Whitney test). (B) Quantification of mitochondrial morphology (either reticular, collapsed, or netted) for RID-Num1 cells is shown as the mean ± SD. Timepoints after addition of rapamycin are shown. Data for WT and Δ*num1* are duplicated from Figure 1C. Each black dot represents the average for one biological replicate, with >100 cells per replicate. Three biological replicates are shown. The black line denotes the grand mean. *p* values are in comparison to WT. * *p* < 0.05; n.s. not significant (Mann-Whitney test). (C) Quantification of mitochondrial morphology (either reticular, collapsed, or netted) for RID^AID^-Num1 cells is shown as the mean ± SD. Timepoints after 30 minute incubation in rapamycin and subsequent addition of auxin are shown. Data for WT and Δ*num1* are duplicated from A. Each black dot represents the average for one biological replicate, with >100 cells per replicate. Three biological replicates are shown. *p* values are in comparison to WT, *p* values in parenthesis are in comparison to Δ*num1*. *** *p* < 0.001; **** *p* < 0.0001; n.s. not significant (Mann-Whitney test).

We also examined mitochondrial morphology in RID-Num1 cells in the *DNM1-GFP* background, which exacerbates the division phenotype in cells lacking Num1. Since RID-Num1 also utilized GFP, we constructed a version that lacked GFP for these experiments. In uRID-Num1 *DNM1-GFP* cells, 35% of cells exhibited a collapsed mitochondrial network and 25% of cells exhibited a net phenotype, similar to Δ*num1 DNM1-GFP* cells (Figure 5B). In iRID-Num1 *DNM1-GFP* cells, near wildtype mitochondrial morphology was restored over time. Thirty minutes after rapamycin addition, the percentage of iRID-Num1 cells with a collapsed mitochondrial network was dramatically reduced and the vast majority of cells exhibited a reticular mitochondrial network (Figure 5B). Interestingly, netted mitochondrial networks persisted (24% of cells) thirty minutes post-rapamycin addition (Figure 5B). However, the percentage of iRID-Num1 cells with netted mitochondrial networks decreased to 19% three hours after rapamycin addition, and further decreased to 12% twenty-one hours after rapamycin addition (Figure 5B). These results suggest that mitochondrial morphology in cells with netted mitochondria takes longer to restore, perhaps because several mitochondrial division events are required to break apart the netted tubules. Eventually, mitochondrial tethering with iRID-Num1 did rescue this division-related morphological defect.

We then used the RID^AID^-Num1 system to measure the kinetics of disruption of mitochondrial division following the loss of mitochondria-PM contact. After three hours in auxin, a time point where most dRID^AID^-Num1 cells lacked Num1 clusters and cortical tethering of mitochondria was dramatically reduced (Figure 3C-D), division rates dropped to levels comparable to uRID-Num1 and Δ*num1* cells (Figure 5C). These results indicate that when mitochondria-PM contact is lost, mitochondrial division is disrupted.

### Num1-mediated mitochondria-plasma membrane tethering is favorable for mitochondrial division

The data presented above indicate that mitochondrial division rates are restored following the formation of Num1-mediated mitochondria tethering sites. However, based on these data, we cannot discern if the role of Num1 in mitochondrial division extends beyond tethering mitochondria to the PM. To test if simply tethering mitochondria to the PM is sufficient to increase rates of mitochondrial division, we engineered an inducible mitochondria-PM tethering system that is independent of Num1. In this system, we expressed the mitochondria-binding domain of Mdv1 (amino acids 1-241, which is also known as the Mdv1 N-terminal extension; Mdv1NTE), as a fusion to FRB-GFP (Figure 6A) (Tieu *et al*., 2002; Cerveny and Jensen, 2003; Zhang and Chan, 2007). FRB-GFP-Mdv1NTE was expressed from an exogenous locus so the native *MDV1* gene was not disrupted. Since Mdv1 is involved in mitochondrial division, we verified that the expression of FRB-GFP-Mdv1NTE in an otherwise wild type background did not affect mitochondrial division (Figure S6) (Tieu *et al*., 2002). To recruit FRB-GFP-Mdv1NTE to the PM in the presence of rapamycin, we co-expressed Pil1 as a fusion to the other component of the rapamycin-induced dimerization system, FKBP12 (Figure 6A). Pil1 is a major component of eisosomes, which are multi-protein assemblies that form discrete stable puncta at the PM (Walther *et al*., 2006). We refer to this artificial mitochondria-PM tether as RID-Mdv1 (Figure 6A). RID-Mdv1 is an inducible version of a constitutive Mdv1NTE-Pil1 artificial tether we used previously to test whether restoring contact or proximity between mitochondria and the PM was able to rescue the respiratory growth defect of Δ*num1* cells (White *et al*., 2022).

**Figure 6:**
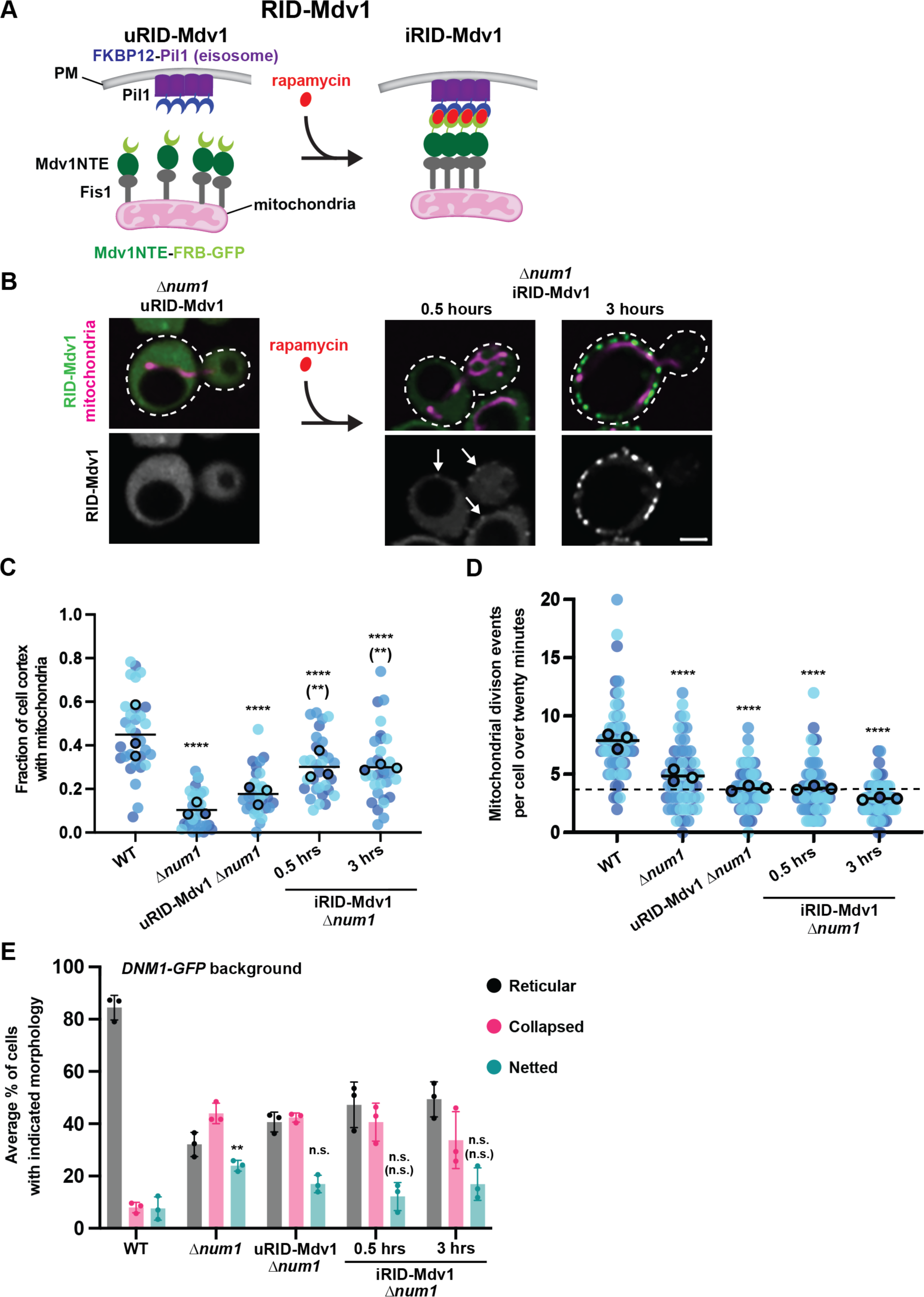
Num1-independent mitochondrial-PM tethering is not favorable for mitochondrial division. (A) Schematic depicting the Num1-independent rapamycin-induced dimerization system using a domain of Mdv1 (RID-Mdv1). The schematic on the left shows the FRB-GFP-Mdv1 and FKBP12-Pil1 constructs before addition of rapamycin and the panel on the right shows the assembled mitochondria-PM tether after rapamycin addition. (B) RID-Mdv1 cells expressing mito-Red were analyzed by fluorescence microscopy. Single mid-slice images are shown. RID-Mdv1 cells without any treatment (uRID-Mdv1) and with rapamycin added (iRID-Mdv1) are shown. A different cell is shown for each condition. The top panel shows a merge of the GFP and mito-Red signals and the bottom panel shows the GFP signal in greyscale, with consistent brightness/contrast. White arrows indicate RID-Mdv1 clusters for iRID-Mdv1 after 30 minutes of rapamycin addition. Bar, 2 µm. (C) Quantification of the fraction of the cell cortex at mid-cell occupied by mitochondria for RID-Mdv1 cells. Each dot represents a single cell, and each biological replicate is represented by a different color. n = 30 cells. *p* values are in comparison to WT, *p* values in parenthesis are in comparison to uRID-Mdv1 Δ*num1*. ** *p* < 0.01; **** *p* < 0.0001 (Mann-Whitney test). (D) Quantification of mitochondrial division events per cell over a twenty-minute time-lapse movie for RID-Mdv1 cells treated with rapamycin for the indicated times. Each dot represents a single cell. There are 20 cells per biological replicate, and the three biological replicates are represented in different colors. The black line denotes the grand mean. *p* values are in comparison to WT. **** *p* < 0.0001 (Mann-Whitney test). (E) Quantification of mitochondrial morphology (either reticular, collapsed, or netted) for RID-Mdv1 cells expressing Dnm1-GFP is shown as the mean ± SD. Timepoints after addition of rapamycin are shown. Each dot represents the average for one biological replicate, with >100 cells per replicate. Three biological replicates are shown. *p* values are in comparison to WT, *p* values in parenthesis are in comparison to uRID-Mdv1 Δ*num1*. ** *p* < 0.01; n.s. not significant (Mann-Whitney test).

We examined whether RID-Mdv1-mediated mitochondria-PM tethering was able to rescue mitochondrial morphology and division rates in cells lacking Num1. In RID-Mdv1 Δ*num1* cells in the absence of rapamycin (uRID-Mdv1), FRB-GFP-Mdv1 was localized diffusely in the cytosol and the mitochondrial division defect observed was similar to that observed in Δ*num1* cells (Figure 6B and D). Following the addition of rapamycin, induced RID-Mdv1 (iRID-Mdv1) clusters on the PM were visible within thirty minutes and became more abundant after three hours (Figure 6B). Consistent with the formation of cortical RID-Mdv1 clusters, cortical mitochondrial tethering was restored to near wild type levels in iRID-Mdv1 cells thirty minutes after rapamycin addition (Figure 6C), similar to iRID-Num1 (Figure 2D). We then assessed division rates in iRID-Mdv1 Δ*num1* cells. In contrast to iRID-Num1 cells, there was no increase in mitochondrial division rates in iRID-Mdv1 Δ*num1* cells (Figure 6D). We also examined mitochondrial morphology in uRID-Mdv1 and iRID-Mdv1 Δ*num1* cells in the *DNM1-GFP* background, which exacerbates the mitochondrial division defect in cells lacking Num1. We found that the percentage of cells that exhibited netted mitochondrial networks did not change following the formation of RID-Mdv1 tethers (Figure 6E), consistent with our finding that division rates were not rescued following RID-Mdv1-mediated mitochondrial-PM tethering (Figure 6D). These results indicate that restoring mitochondria-PM contact with the RID-Mdv1 tether is not sufficient to rescue the mitochondrial division defect observed in the absence of Num1.

## DISCUSSION

One of the many functions ascribed to Num1 is a role in mitochondrial division (Cerveny *et al*., 2007). Here, we further dissect this role and provide evidence that Num1 is not required for mitochondrial division, consistent with our previous study (Lackner *et al*., 2013), but does enhance mitochondrial division. To investigate the relationship between Num1-mediated mitochondrial tethering and mitochondrial division, we designed a regulatable version of the native Num1 tether, called RID-Num1. While regulatable artificial membrane tethers have been used to study MCS functions (Csordás *et al*., 2010; Booth *et al*., 2016, 2021; Kwak *et al*., 2020; Benedetti, 2021; Katona *et al*., 2022), this is the first use of the rapamycin-inducible dimerization system to reconstitute a native MCS. By combining the rapamycin-inducible dimerization system with the auxin-inducible degradation system, we developed a system called RID^AID^-Num1, in which we were able to establish and deplete native Num1-mediated mitochondrial tethering sites. We found that depletion of Num1-mediated contact sites results in a reduction of mitochondrial division rates, and that mitochondrial division rates are increased, approaching wild type levels, shortly after the reestablishment of Num1-mediated mitochondrial tethering. These results support the idea that Num1-mediated mitochondria-PM tethering facilitates mitochondrial division. To determine if the role of Num1 in mitochondrial division extends beyond simply tethering mitochondria to the PM, we developed a regulatable tethering system, RID-Mdv1, to increase mitochondria-PM contact in a Num1-independent manner. We found that even though RID-Mdv1 restored cortical mitochondria tethering in the absence of Num1, mitochondrial division rates were not restored. Thus, there is a specific feature of Num1-mediated mitochondria-PM tethering that facilitates mitochondrial division.

One possible explanation for these results is that Num1-mediated tethering exerts an ideal amount of tension across the mitochondrial membrane for mitochondrial division to occur. Several studies have found a connection between mitochondrial membrane tension and membrane division (Roux *et al*., 2006; Mahecic *et al*., 2021). There are, on average, 3-4 Num1-mediated mitochondrial tethering sites per cell that are spaced out along the cell cortex (Figure 1 B and C; Kraft and Lackner, 2017). It is possible that the placement and number of Num1 clusters creates an ideal tethering geometry to facilitate mitochondrial division. While RID-Mdv1 increases mitochondria-PM contact, the number and placement of the tethering sites differ from Num1 tethering sites, which may explain why mitochondrial division rates were not restored in the iRID-Mdv1 cells.

A second possibility is that fully reconstituted Num1 regulates mitochondrial division factors, such as Dnm1. It is clear from experiments comparing uRID-Num1 and iRID-Num1 that the fully reconstituted Num1 has a positive impact on mitochondrial division. There is a population of Dnm1 that colocalizes with cortical Num1 clusters overtime (Cerveny *et al*., 2007; Lackner *et al*., 2013). While this population of Dnm1 does not appear to be actively participating in mitochondrial division (Lackner *et al*., 2013), there is likely a functional purpose for the colocalization of Dnm1 with Num1 clusters. It is possible that Num1 could be recruiting and holding Dnm1 in close proximity to the mitochondrial membrane, acting like a reservoir for Dnm1 so it is accessible when needed. Alternatively, Dnm1 may be playing a division-independent role at this tripartite membrane contact site.

The regulatable MCS systems engineered for this study have broad applications for studying MCS biology and the spatial and temporal relationship between the formation of a MCS and the functions attributed to that MCS. Historically, deletions or mutations of contact site proteins have been used to study MCS functions. Regulatable contact sites are advantageous to these non-conditional mutants; they provide the ability to deplete and re-establish MCSs and observe the downstream consequences in real time, which proved valuable in examining the relationship between Num1-mediated mitochondrial tethering and mitochondrial division. The utility of the regulatable native and artificial tethers described here extend beyonds this study. We have employed RID-Num1 and RID-Mdv1 to assess a novel connection between MECA and the distribution of lipid species in the PM (Casler et al. in prep). It is likely possible to expand the use of RID-Num1 and RID^AID^-Num1 to mammalian cells. Since Num1 recognizes conserved features of the mitochondrial and plasma membranes (Yu *et al*., 2004; Ping *et al*., 2016; Kraft and Lackner, 2019), it is possible that the regulatable Num1 tether can be used as an exogenous system in mammalian cells to control mitochondrial positioning relative to the PM. Given the rapidly growing field of MCS biology and the functional complexity of the proteins that form MCSs, tools such as those described here will be critical for diving deeper into the mechanisms underlying the cellular processes facilitated by MCSs.

## MATERIALS AND METHODS

### Strains and plasmids

Supplemental Tables 1 and 2 list all strains and primers used for this study, respectively. Strains were constructed using standard yeast techniques (PCR-based targeted homologous recombination and transformation). Primer sets used to amplify deletion cassettes are named F1 and R1 and primer sets used to amplify C-terminal gene tagging cassettes are named F5 and R3 (Longtine *et al*., 1998; Janke *et al*., 2004; Sheff and Thorn, 2004). Plasmids used in this study are listed in Supplemental Table 3. All strains and plasmids will be made available upon request.

### Imaging

For Figure 1B, 1C and 1E, all strains were grown to mid-log phase in synthetic complete +2% (wt/vol) dextrose media with 2x adenine (SCD). Cells were imaged on an agarose pad on a depression slide. Single time point images were captured on a Nikon Spinning Disk Confocal System fitted with a CSU-W1 dual-disk spinning disk unit (Yokogawa). A 60X (NA-1.42) objective and a Hamamatsu ORCA Fusion Digital CMOS camera were used for image capture. Images were captured and deconvolved with Elements software (Nikon).

For Figure 1D, all strains were grown to mid-log phase in synthetic complete+2% (wt/vol) dextrose media with 2x adenine (SCD). Cells were imaged on concanavalin A-treated dishes (2 mg/mL in water). Twenty-minute time-lapse movies were captured on a Nikon Spinning Disk.

For Figure 2B-D, all strains were grown to mid-log phase in synthetic complete+2% (wt/vol) dextrose media with 2x adenine (SCD). A final concentration of 1 µM rapamycin was added to some of the cultures for various lengths of time, as indicated in the figure. Cells were imaged on an agarose pad on a depression slide. Single time point images were captured on a Leica Spinning Disk Confocal System fitted with a CSU-X1 spinning-disk head (Yokogawa). A PLAN APO 100X and an Evolve 512 Delta EMCCD camera (Photometrics) were used for image capture. Images were captured with Metamorph (Molecular Devices) and deconvolved using AutoQuant X3’s (Media Cybernetics) iterative, constrained, three-dimensional deconvolution method. Linear adjustments to brightness and contrast were performed using FIJI and Photoshop (Adobe). Deconvolved images are shown in the figure.

For Figure 3B-D, all strains were grown to mid-log phase in synthetic complete+2% (wt/vol) dextrose media with 2x adenine (SCD). A final concentration of 1 µM rapamycin was added to cells notated as iRID and incubated for 30 minutes before imaging. For dRID cultures, 1 µM rapamycin was added for 30 minutes, and then 1 mM auxin at pH 6.4 was added for the indicated time before imaging. Cells were imaged on an agarose pad on a depression slide. Single time point images were captured as described for Figure 2B.

For Figure 4A, all strains were grown to mid-log phase in synthetic complete+2% (wt/vol) dextrose media with 2x adenine (SCD). Cells were imaged on an agarose pad on a depression slide. Single time point images were captured on Leica TCS SP8 LSCM. A 63X (NA-1.4) objective and HyD detectors were used for image capture. Huygens software was used for image deconvolution.

For Figure 5A and 5C, all strains were grown to mid-log phase in synthetic complete+2% (wt/vol) dextrose media with 2x adenine (SCD). A final concentration of 1 µM rapamycin was added to cells notated as iRID and incubated for the indicated times before imaging. For dRID cultures, 1 µM rapamycin was added for 21 hours, and then 1 mM Auxin at pH 6.4 was added for three hours before imaging. Cells were imaged on concanavalin A-treated dishes (2 mg/mL in water). Twenty-minute time-lapse movies with 30 second timepoints were captured on Nikon Sora as described for Figure 1B.

For Figure 5B, all strains were grown to mid-log phase in synthetic complete+2% (wt/vol) dextrose media with 2x adenine (SCD). Cells were imaged on an agarose pad on a depression slide. Single time point images were captured on Nikon Sora as described for Figure 1B.

For Figure 6B and D, all strains were grown to mid-log phase in synthetic complete +2% (wt/vol) dextrose media with 2x adenine (SCD). Cells were imaged on an agarose pad on a depression slide. Single time point images were captured on Nikon Sora as described for Figure 1B.

### Image Quantifications

For mitochondria morphology quantifications, single time point maximum projection images were analyzed on a per cell basis; each cell’s mitochondrial network was classified as reticular, collapsed, or netted. See Supplemental Figure 1 for examples of each phenotype.

For mitochondrial division event quantifications, a twenty-minute time-lapse maximum projection movie was analyzed and events of mitochondrial division were counted on a per cell basis.

For Num1-GFP, RID-Num1, and RID-AID cluster quantifications, single time point maximum projection images were analyzed for the number of GFP foci.

For mitochondria-plasma membrane contact quantifications, the proportion of the cell cortex in which mitochondrial signal was present was quantified similar to Kraft and Lackner (Kraft and Lackner, 2019). A five-pixel wide line was drawn around the circumference of the cell in FIJI software. The intensity values of the mitochondria channel were collected along the drawn line. Background intensity values were determined by drawing a 5 µM long, five-pixel wide line in the background for each field of view used. The number of data points above the average background intensity value for the corresponding field of view divided by the total number of data points was plotted.

For RID-Num1 and RID-AID cluster ER association quantifications, single time point single-slice images were used to score if a GFP foci overlapped with ER signal.

### Western blots

Cells were grown to mid-log in yeast extract/peptone with 2% (w/v) dextrose (YPD), with 1 µM rapamycin and/or 1 mM auxin added for the time indicated in the figure. Cells were harvested by centrifugation (3000 x g for 1 minute) and whole-cell extracts were prepared by a NaOH lysis and trichloroacetic acid precipitation procedure. Pellets were resuspended in 50 µL of MURB (100 mM MES, pH 7, 1% SDS, and 3 M urea). Whole cell extracts were analyzed by SDS-PAGE and Western Blot using Revert Total Protein Stain (LI-COR Biosciences) as a control. Antibodies used were anti-GFP (Abclonal Rabbit anti-GFP-Tag pAb 1:2,000) and anti-FKBP12 (Abcam Rabbit anti-FKBP12 ab2918 1:1,000) as primaries and goat anti-rabbit IgG DyLight 800 (Thermo Fisher Scientific 1:15,000) as a secondary antibody. The total protein stain and immunoreactive bands were detected with the Odyssey Infrared Imaging System (LI-COR Biosciences). Western blots were quantified using ImageStudio (LI-COR Biosciences) by normalizing the intensity of the immunoreactive band of interest to the total protein for each sample.

### Growth assays

To analyze growth by serial dilution, cells were grown in YPD medium overnight at 30°C. Cells were diluted and allowed to grow to mid-log at 30°C, then 0.2 OD600 of cells were pelleted (3000 x g for 1 minute) and resuspended in water to a final OD600 of 0.5. Fivefold serial dilutions were conducted, spotted onto YPD or YPEG agar plates, and grown at 37°C as indicated.

To analyze growth quantitatively by colony size, cells were grown as described above to achieve mid-log growth. Analysis was performed as described in White et al. (White *et al*., 2022). Two 1:100 serial dilutions were performed to achieve cell concentrations of ∼100 cells/ml. 150 μl of the diluted cell culture were plated onto both YPD and YPEG plates to achieve roughly 100 individual colonies per plate and left to grow at 35°C. YPD plates were removed from the incubator after 48 h of growth, and YPEG plates were removed from the incubator after 120 h of growth. For analysis, plates were scanned and images were analyzed using OpenCFU software to identify the radius (in pixels) of individual colonies as a measure of colony size (Geissmann, 2013). In the analysis, colonies were excluded if: (1) the box that defined the area measured for a particular colony encompassed more than one colony; (2) the box that defined the area measured did not completely encompass the colony being measured; or (3) if colony morphology was significantly impacted by touching the side of the plate or other colonies, such that area being measured was not accurate. All strains were normalized to the average colony size of the wild type sample grown in the same condition on the same day. Specifically, for each round of the assay, the average radius size of wild type colonies was determined for each condition. Colony size for all of the strains from the same experimental round and condition was divided by the average radius size of wild type colonies. For graphs that include data collated from experiments that were each normalized to their own wild type data set, the wild type data sets from all experiments are included in the graph. In addition to showing the grand (i.e. overall pooled) mean of the single colony data, the mean of each independent experiment is shown using circles. Statistical analyses were performed with R programs using a two-tailed unpaired t-test with a 95% confidence interval. The independent experimental means for each genotype were used for the statistical comparisons as described in Lord et al. (Lord *et al*., 2020). Designations of significance are indicated in the figure legends.

## ACKNOWLEDGEMENTS

We thank the past and present Lackner lab members for scientific discussions, ideas, and helpful feedback. Special thanks to Erica Rosario for assisting with data visualization and Kaylee Gonzalez for assisting with yeast strain construction. We also thank Northwestern’s Cell Biology Supergroup, the Wignall-Lackner (WiLa) Cell Biology Group, and the Biotechnology Training Program for suggestions and feedback. Thank you to the Brickner lab for providing yeast strains and plasmids. All microscopy was performed at the Biological Imaging Facility at Northwestern University (RRID:SCR_017767), supported by the Chemistry for Life Processes Institute, the Northwestern University Office for Research, the Department of Molecular Biosciences, and the Rice Foundation. Special thanks to Jessica Hornick and Tong Zhang for assistance with microscopy. C.S.H. was supported by National Institutes of Health Biotechnology Training Program 5T32GM008449-25 and National Science Foundation Graduate Research Fellowship Program under Grant No. DGE-1842165. J.C.C is supported by National Institutes of Health, National Institute of General Medical Sciences grant 1F32GM145160-01. L.L.L. is supported by National Institutes of Health, National Institute of General Medicine Sciences grant R01GM120303.

