## Supplemental Data for "Temporal control of Num1 contact site formation reveals a relationship between mitochondrial division and mitochondria-plasma membrane tethering"

Clare S. Harper, Jason C. Casler, and Laura L. Lackner*

Department of Molecular Biosciences, Northwestern University, Evanston, IL, 60208, USA.

**Supplementary Figures**

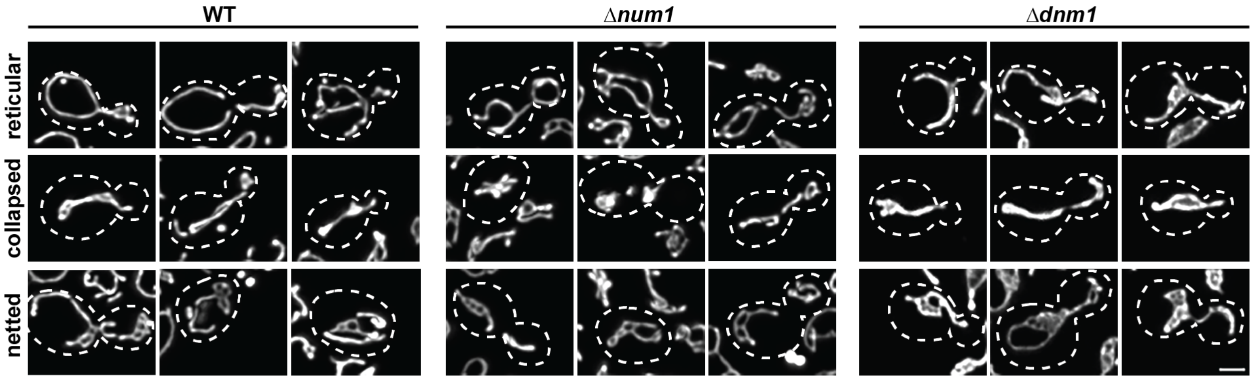

**Figure S1: Mitochondrial morphologies across Δ*num1* and Δ*dnm1* mutants.** Wildtype, Δ*num1*, and Δ*dnm1* cells expressing mito-Red were analyzed by fluorescence microscopy. Maximum projection images are shown. Three representative cells are shown for each mitochondrial morphology phenotype, with consistent brightness/contrast. Bar, 2 μm.

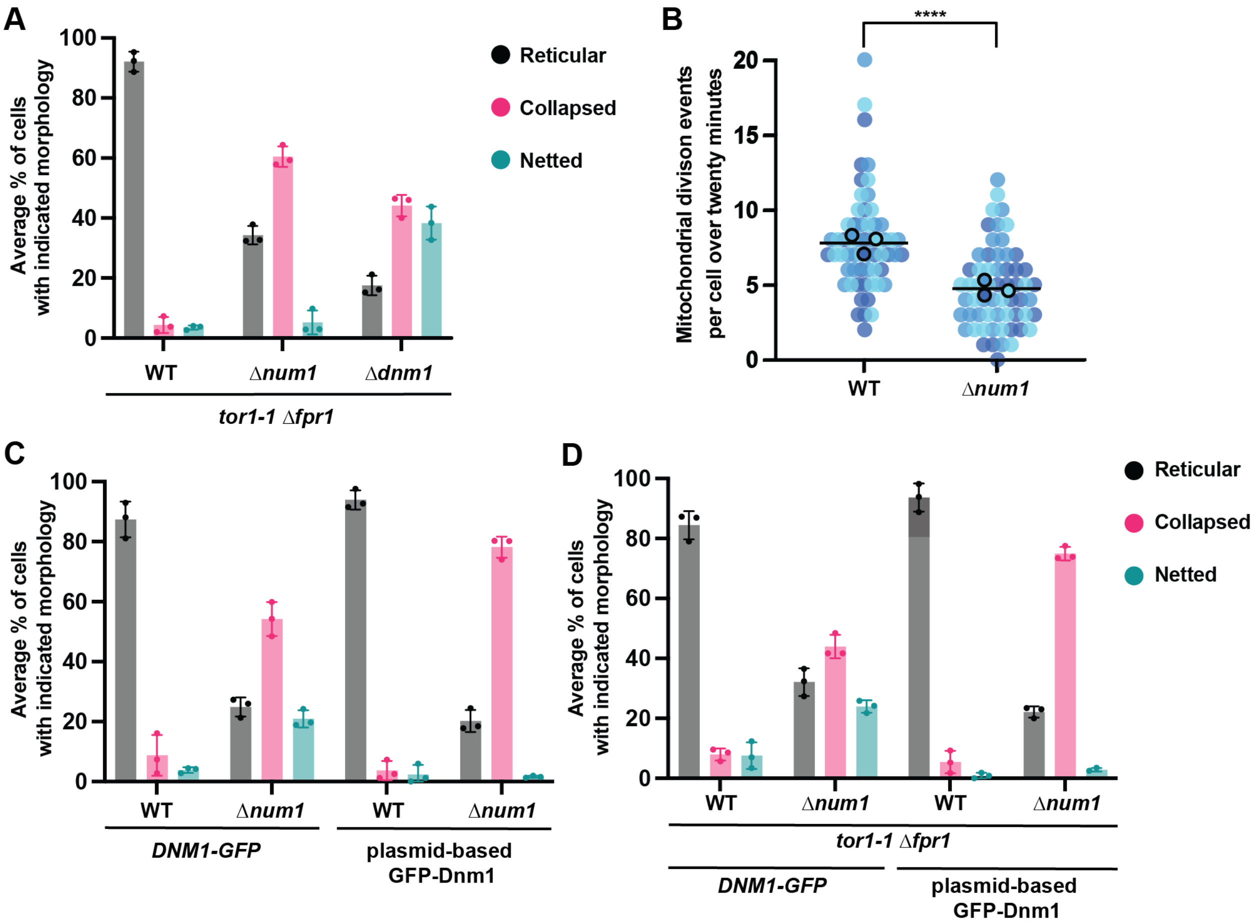

**Figure S2: Rapamycin resistant strain shows similar defects in mitochondrial division upon loss of Num1 or Dnm1 compared to W303.** (A) Quantification of mitochondrial morphology (either reticular, collapsed, or netted) for rapamycin resistant WT, Δ*num1*, and Δ*dnm1* cells is shown as the mean ± SD. Each dot represents the average for one biological replicate, with >100 cells per replicate. Three biological replicates are shown. (B) Quantification of mitochondrial division events per cell over a twenty-minute time-lapse movie for rapamycin resistant WT and Δ*num1* cells. Each dot represents a single cell, and each outlined dot represents the average of a given replicate. There are 20 cells per biological replicate, and the three biological replicates are represented in different colors. The black line denotes the grand mean of the three replicate averages. *p* values are in comparison to WT. **** $p < 0.0001$ (Mann-Whitney test). (C) Quantification of mitochondrial morphology (either reticular, collapsed, or netted) for WT and

Δ*num1* cells expressing Dnm1-GFP or plasmid-based GFP-Dnm1 is shown as the mean ± SD. Each dot represents the average for one biological replicate, with >100 cells per replicate. Three biological replicates are shown. Data for WT and Δ*num1* with *DNM1-GFP* duplicated from Figure 1C. (D) Quantification of mitochondrial morphology (either reticular, collapsed, or netted) for rapamycin resistant WT, Δ*num1*, and Δ*dnm1* cells expressing either Dnm1-GFP or plasmid-based GFP-Dnm1 is shown as the mean ± SD. Each dot represents the average for one biological replicate, with >100 cells per replicate. Three biological replicates are shown.

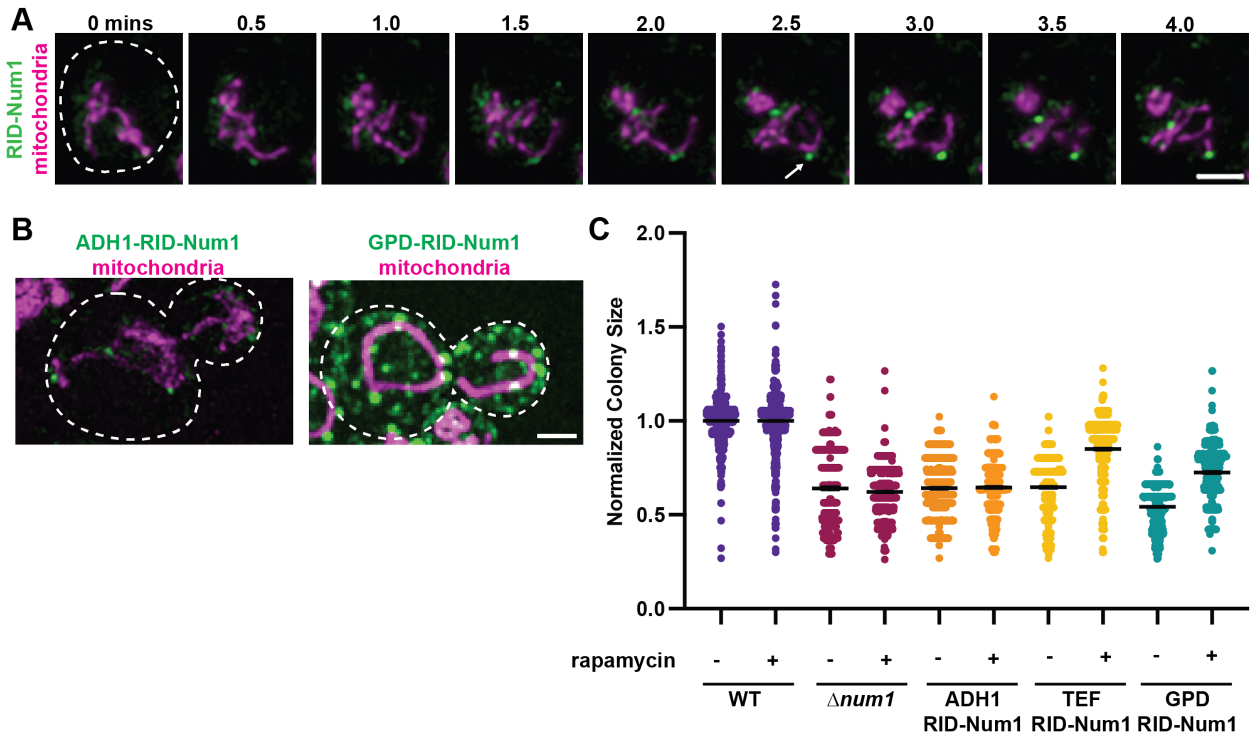

**Figure S3: RID cluster formation and validating promoter choice for expressing FKBP12-Num1PH for the RID-Num1 system.** (A) RID-Num1 cells expressing mito-Red were analyzed by live-cell time-lapse fluorescence microscopy. Cells were imaged in a dish with rapamycin media flowed in during the movie. Arrow indicates a newly formed RID-Num1 cluster. Maximum projection images from 0.5 minute timepoints are shown. Bar, 2 µm. Timepoints are from Video 1. (B) Wildtype, Δ*num1*, and RID-Num1 cells (with FKBP12-Num1PH under the ADH or GPD promoter) expressing mito-Red were analyzed by fluorescence microscopy. Maximum projection images are shown. Bar, 2 µm. (C) Quantification of WT, Δ*num1*, and RID-Num1 (with FKBP12-Num1PH under the ADH, TEF, or GPD promoters) colony size, with cells grown at 35°C in respiratory growth conditions (YPEG solid media) for 5 days. Rapamycin at 1µM was added to plates, as indicated. The graph is a violin plot of the radius (in pixels) of colonies normalized to the mean radius of WT colonies for the respective experiment. The black line denotes the grand mean of at least three independent experiments. n>300 colonies.

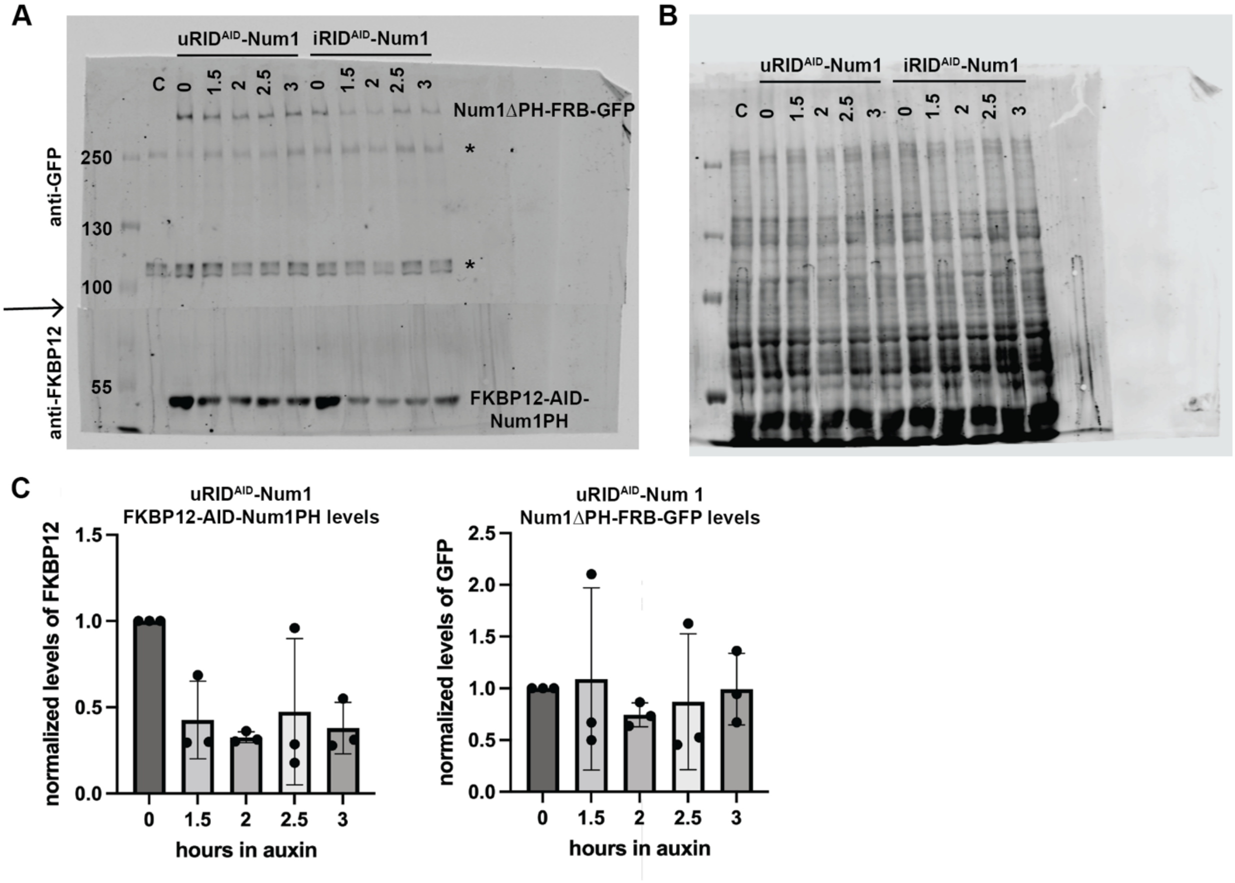

**Figure S4: Western blot and total protein stain for RID$^{AID}$-Num1.** (A) Western blot of RID$^{AID}$-Num1 components (detecting Num1∆PH-FRB-GFP with an α-GFP antibody on the top half of the blot and FKBP12-Num1PH with an α-FKBP12 antibody on the bottom half of the blot, separated as indicated by the arrow). iRID$^{AID}$-Num1 cells were incubated with rapamycin for 30 minutes and uRID$^{AID}$-Num1 cells had no rapamycin treatment. Auxin was then added, and the cells were analyzed at the indicated times (in hours) after auxin addition. "C" indicates a WT control. Asterisks denote non-specific bands from the α-GFP antibody. (B) Total protein stain of blot in A (as described in Methods). (C) Quantification of normalized protein levels is shown as the mean ± SD. Each dot indicates an independent experiment. n=3.

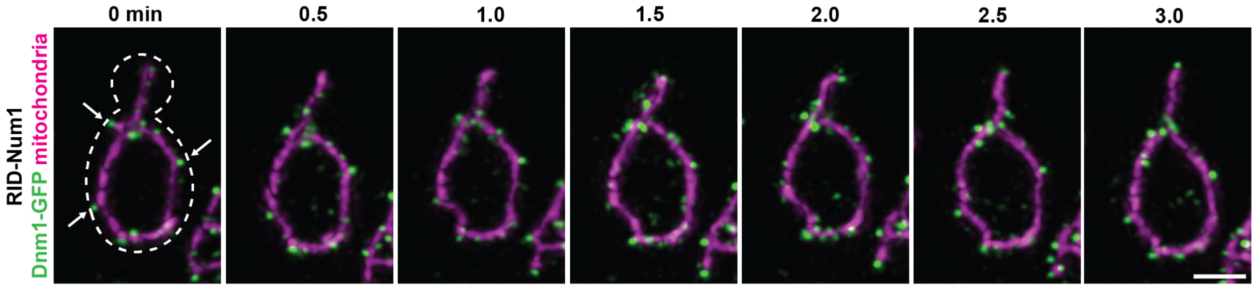

**Figure S5: Cortical Dnm1 foci are present in the RID-Num1 strain, similar to WT.** RID-Num1 cells expressing *DNM1-GFP* and mito-RED were analyzed by live-cell time-lapse fluorescence microscopy. Arrows indicate cortical Dnm1-GFP foci. Maximum projection images from 0.5 minute timepoints are shown. Bar, 2 μm. Timepoints are from Video 2.

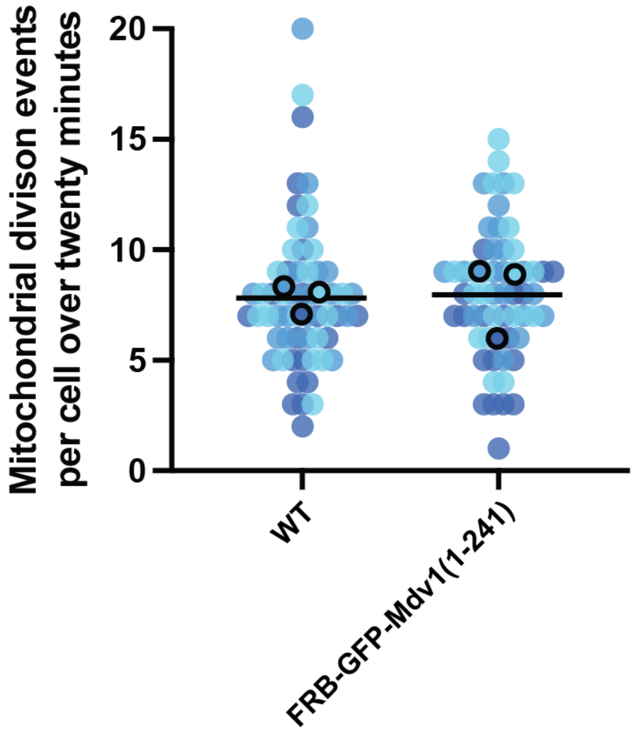

**Figure S6: GPD::FRB-GFP-Mdv1(1-241) construct for RID-Mdv1 strain does not impair mitochondrial division.** Quantification of mitochondrial division events per cell over a twenty-minute time-lapse movie for cells expressing the GPD::FRB-GFP-Mdv1(1-241) plasmid (used in the RID-Mdv1 strain) and mito-Red. Each dot represents a single cell, and each outlined dot represents the average of a given replicate. There are 20 cells per biological replicate, and the three biological replicates are represented in different colors. The black line denotes the grand mean of the three replicate averages.

**Video 1: RID-Num1 forms stable clusters that tether mitochondria overtime upon rapamycin addition.** RID-Num1 cells expressing mito-Red were analyzed by live-cell time-lapse fluorescence microscopy. Cells were imaged in a dish with rapamycin media flowed in during the movie. Movie duration is four minutes with 0.5 minute timepoints. Maximum projection is shown. Timepoints from the movie are shown in Figure S3A.

**Video 2: Cortical Dnm1 foci are present in RID-Num1.** iRID-Num1 cells expressing DNM1-GFP and mito-Red were analyzed by live-cell time-lapse fluorescence microscopy. Movie duration is ten minutes with 0.5 minute timepoints. Maximum projection is shown. Timepoints from the movie are shown in Figure S5.

**Supplemental Table 1**

| Strain number (LLY) | Strain name | Genotype | Source |
|---|---|---|---|
| 92 | WT | W303 (ade2–1;leu2–3;his3–11,15;trp1–1;ura3–1;can1–100) | |
| 3351 | | W303 mitoRed::LEU/NAT | White et al 2022 |
| 3352 | Δ*num1* | W303 Δ*num1*::KAN mitoRed::LEU/NAT | White et al 2022 |
| 5021 | Δ*dnm1* | W303 Δ*dnm1*::NAT mitoRed::LEU/NAT | White et al 2022 |
| 3242 | *DNM1-GFP* | W303 *DNM1-yEGFP*::HIS mitoRed::LEU/NAT | |
| 5668 | | W303 Δ*num1*::KAN *DNM1-yEGFP*::HIS mitoRed::LEU/NAT | |
| 2957 | Rapamycin resistant WT | W303 *TOR1-1* Δ*fpr1*::NAT | Haruki et al 2008 Gruber et al 2006 Gift from Jason Brickner. |
| 5634 | | W303*TOR1-1* Δ*fpr1*::NAT mitoRed::LEU/NAT | |
| 5325 | Rapamycin resistant | W303 *TOR1-1* Δ*fpr1*::NAT Δ*num1*::KAN mitoRED::LEU/NAT | |

| | | |
|---|---|---|
| | *Δnum1* | |
| | Rapamycin resistant *Δdnm1* | W303 *TOR1-1 Δfpr1*::NAT *Δdnm1*::KAN mitoRED::LEU/NAT |
| 5691 | Rapamycin resistant *DNM1-GFP* | W303 *TOR1-1 Δfpr1*::NAT *DNM1-yEGFP*::HIS mitoRed::LEU/NAT |
| 5667 | | W303 *TOR1-1 Δfpr1*::NAT *Δnum1*::KAN *DNM1-yEGFP*::HIS mitoRed::LEU/NAT |
| 5635 | RID-Num1 | W303 *TOR1-1 Δfpr1*::NAT *NUM1ΔPH-FRB-GFP*::HIS *TEF-FKBP12-NUM1(PH domain)*::URA mitoRed::LEU/NAT |
| 5102 | RID-Num1 | W303 *TOR1-1 Δfpr1*::NAT *TEF-FKBP12-AID-NUM1(PH domain)*::TRP *NUM1ΔPH-FRB-GFP*::HIS *TIR1*::URA mitoRed::LEU/NAT |
| 4932 | | W303 *TOR1-1 Δfpr1*::NAT *NUM1-GFP*::KAN mitoRED::LEU/NAT |
| 5080 | | W303 *TOR1-1 Δfpr1*::NAT *NUM1ΔPH-FRB-GFP*::HIS *TEF-FKBP12-NUM1(PH domain)*::URA *PHO88-mCherry*::LEU |

| | | |
|---|---|---|
| 5103 | | W303 *TOR1-1* Δ*fpr1*::NAT *NUM1ΔPH-FRB-GFP*::HIS *TEF-FKBP12-NUM1(PH domain)*::URA *PHO88-mCherry*::LEU Δ*lnp1*::KAN |
| 5513 | | W303 *TOR1-1* Δ*fpr1*::NAT *PHO88-mCherry*::LEU *NUM1-GFP*::HIS |
| 5514 | | W303 *TOR1-1* Δ*fpr1*::NAT Δ*lnp1*::KAN *PHO88-mCherry*::LEU *NUM1-GFP*::HIS |
| 5553 | | W303 *TOR1-1* Δ*fpr1*::NAT *NUM1ΔPH-FRB*::HIS *TEF-FKBP12-NUM1(PH domain)*::URA mitoRED::LEU/NAT *DNM1-yEGFP*::HIS |
| 5102 | RID$^{AID}$-Num1 | W303 *TOR1-1* Δ*fpr1*::NAT *TEF-FKBP12-AID-NUM1(PH domain)*::TRP *NUM1ΔPH-FRB-GFP*::HIS *TIR1*::URA mitoRed::LEU/NAT |
| 5827 | RID-Mdv1 | W303 *TOR1-1* Δ*fpr1*::NAT *PIL1-FKBP12*::KAN *GPD-FRB-yEGFP-MDV1NTE*::URA Δ*num1*::KAN mitoRED::LEU/NAT |
| 5943 | | W303 *TOR1-1* Δ*fpr1*::NAT *PIL1-FKBP12*::KAN *GPD-FRB-yEGFP-* |

| | | |
|---|---|---|
| | | *MDV1NTE*::URA Δ*num1*::KAN mitoRED::LEU/NAT *DNM1-yEGFP*::HIS |
| 4577 | | W303 *TOR1-1* Δ*fpr1*::NAT Δ*kar9*::NAT |
| 4578 | | W303 *TOR1-1* Δ*fpr1*::NAT *NUM1ΔPH-FRB-GFP*::HIS Δ*kar9*::NAT |
| 5470 | | W303 *TOR1-1* Δ*fpr1*::NAT *NUM1ΔPH-FRB-GFP*::HIS *TEF-FKBP12-Num1(PH domain)*::URA Δ*kar9*::NAT |
| 5927 | Mdv1 construct control | W303 *TOR1-1* Δ*fpr1*::NAT *GPD-FRB-yEGFP-Mdv1NTE*::URA mitoRED::LEU/NAT |

**Supplemental Table 2**

| **Oligo number** | **Oligo name** | **Oligo sequence** |
|---|---|---|
| 174 | Num1 F1 | CGAGTAAAGACGCAACGGTCAAGGCTTTCCACGAGACGTTCGAATCGGATCCCCGGGTTAATTAA |
| 175 | Num1 R1 | GATTATTATTGTTCTTAATTTACTTAGAGTTATTTAGTTTTTTTAAGAATTCGAGCTCGTTTAAAC |
| 125 | Dnm1 F1 | GAGTTTATCATTAAGTAGCTACCAGCGAATCTAAATACGACGGATAAAGACGGATCCCCGGGTTAATTAA |

| | | |
|---|---|---|
| 126 | Dnm1 R1 | GCCCGCAATGTTGAAGTAAGATCAAAAATGAGATGAATTATGCAAGAATTCGAGCTCGTTTAAAC |
| 463 | Dnm1 F5 | GAAATCACTCGGAGTTTATAAAAAGGCTGCAACCCTTATTAGTAATATTCTGGGTGACGGTGCTGGTTTA |
| 464 | Dnm1 R3 | CTATAATCACGCCCGCAATGTTGAAGTAAGATCAAAAATGAGATGAATTATGCAATCGATGAATTCGAGCTCG |
| 259 | Num1∆PH F5 | CTGCCAGTAGGACGGCATCTTTACAAACTTTAGCATCATTGGGTGACGGTGCTGGTTTA |
| 178 | Num1 R3 | GATTATTATTGTTCTTAATTTACTTAGAGTTATTTAGTTTTTTTAATCGATGAATTCGAGCTCG |
| 177 | Num1 F5 | GACATAGAGTACCACAAAGCCGATCATTTGGCAATTTACGAGGTGACGGTGCTGGTTTA |
| 877 | Lnp1 F1 | CATACAAAGAGGAGATCGGATATAAAAGAATAACATAAATCGGATCCCCGGGTTAATTAA |
| 878 | Lnp1 R1 | GATATAAAAATATATTATATAGGGGTACGTAGTTATTCTAACGCAAAAGAATTCGAGCTCGTTTAAAC |
| 148 | Pil1 F5 | GTCGGACACCAGCAAAGTGAGTCTCTTCCCCAACAAACAACAGCTGGTGACGGTGCTGGTTTA |
| 149 | Pil1 R3 | CTGCTGGTTTTTTTTTTTTTTGTTTCTAATAGATTGTTGATTTATTTTGATCGATGAATTCGAGCTCG |

| | | |
|---|---|---|
| 409 | Kar9 F1 | GTGGCCGCACGAATCTTTGTCTGTAACAGCCTTAAAGAT TTCAGTAGCACTGCCCGGATCCCCGGGTTAATTAA |
| 410 | Kar9 R1 | GAGGGTGAGAGGGAGGATATATAAAAATGTATAAGTATA CAGTTTTAGGTTAGTAGAATTCGAGCTCGTTTAAAC |

**Supplemental Table 3**

| **Plasmid number** | **Plasmid construct** |
|---|---|
| 765 | FRB-GFP |
| 864 | ADH1::FKBP12 x2- Num1 PH domain |
| 863 | TEF::FKBP12 x2- Num1 PH domain |
| 791 | GPD::FKBP12 x2- Num1 PH domain |
| 19 | mitoRED |
| 895 | GPD::FRB-yEGFP-Mdv1NTE |
| 899 | GPD::FRB-Mdv1NTE |
| 400 | ADH1::OsTIR1-9Myc |
| 878 | TEF::FKBP12 x2-AID-Num1 PH domain |
| 702 | FKBP12 X2 |
| 54 | pFA6a–link–yEGFP |

| | |
|---|---|
| 169 | PromDnm1::Dnm1-GFP |
